# m^6^A RNA methylation regulates the fate of endogenous retroviruses

**DOI:** 10.1101/2020.03.24.005488

**Authors:** Tomasz Chelmicki, Emeline Roger, Aurélie Teissandier, Sofia Rucli, François Dossin, Mathilde Dura, Camille Fouassier, Sonia Lameiras, Deborah Bourc’his

## Abstract

Endogenous retroviruses (ERVs) are abundant and heterogenous groups of integrated retroviral sequences that impact genome regulation and cell physiology throughout their RNA-centered life cycle^1^. Failure to repress ERVs is associated with cancer, infertility, senescence and neurodegenerative diseases^2–4^. Here, using an unbiased genome-scale CRISPR knockout screen in mouse embryonic stem cells, we identify m^6^A RNA methylation as a novel means of ERV restriction. Methylation of ERV mRNAs is catalyzed by the complex of methyltransferase-like METTL3/METTL14^5^ proteins whose depletion, along with their accessory subunits, WTAP and ZC3H13, led to increased mRNA abundance of Intracisternal A-particles (IAPs) and related ERVK elements specifically, by targeting their 5’UTR region. Using controlled auxin-dependent degradation of the METTL3/METTL14 enzymatic complex, we showed that IAP mRNA and protein abundance is dynamically and inversely correlated with m^6^A catalysis. By monitoring mRNA degradation rates upon METTL3/14 double degron, we further proved that m^6^A methylation destabilizes IAP transcripts. Finally, similarly to m^6^A writers, triple knockout of the m^6^A readers YTHDF1, DF2 and DF3^6^ increased IAP mRNA abundance. This study sheds light onto a novel function of RNA methylation in protecting cellular integrity by clearing reactive ERV-derived RNA species, which may be especially important when transcriptional silencing is less stringent.

## Main text

Mammalian genomes host millions of copies of retrotransposons, including endogenous retroviruses (ERVs) that derive from past retroviral infections and have integrated as permanent genetic residents. Over the course of evolution, ERVs have invaded mammalian genomes in successive waves and have multiplied and diversified, providing a fertile ground for genomic innovations. However, ERVs potentially compromise genomic integrity by disrupting gene structure and expression, and by triggering chromosome rearrangements^7^. In laboratory mice, roughly 12% of all pathological mutations result from ERV integration events, half of which emanate from a single family of the ERVK class, the Intracisternal A particles (IAP), that comprise ∼2800 full-length copies^8^. In contrast, human-specific ERVs are mostly transposition-defective^8^. However, by providing *cis*-regulatory modules, they can divert regulatory networks and alter cellular states. Moreover, ERVs generate RNA, cDNA, RNA:DNA hybrid species and proteins, the accumulation of which is associated with and may contribute to senescence, cancer and neurodegenerative diseases^9^.

In normal conditions, homeostatic ERV regulation is achieved through multilayered surveillance at different steps of the ERV life cycle. In particular, chromatin-based silencing by DNA methylation and histone modifications and post-transcriptional control through RNA editing and RNA interference (RNAi) have been extensively characterized^2^. However, these control mechanisms are not active in all cell types or developmental periods, suggesting that other ERV limiting pathways have yet to be uncovered.

To identify novel factors participating in ERV control, we carried out a CRISPR-Cas9 loss-of-function screen for a highly active mouse ERV representative: the IAPEz family. We engineered mouse embryonic stem cells (ESCs) to carry a constitutively expressed Cas9 transgene at the *TIGRE* locus, and a reporter cassette with IAPEz regulatory elements inserted at the *ROSA26* locus: IAPEz(5’LTR + 5’UTR + 3-60n *gag*)-GFP-Blast^R^ reporter (Supplementary Table 1), in which LTR stands for long terminal repeat, UTR for untranslated region and Blast^R^ indicates resistance to blasticidin (Fig. 1a, Extended Data Fig. 1a). Endogenous IAPEz 5’LTR+5’UTR promoter sequences are globally methylated and silenced in ESCs although basal IAP mRNA levels can be detected^10^. Placing a doxycycline (dox)-responsive promoter upstream of the LTR sequence allowed us to test reporter reactivation upon dox induction (Extended Data Fig. 1b,c) and to adjust minimal blasticidin concentration for selection. We also showed that the IAP reporter responds in the absence of known IAP repressors, by transducing cells with single guide (sg)RNAs against the KRAB-associated protein 1 (KAP1)^11^ (Extended Data Fig. 1d,e, Supplementary Fig. 1, Supplementary Table 2, Supplementary Table 3).

**Fig. 1.**
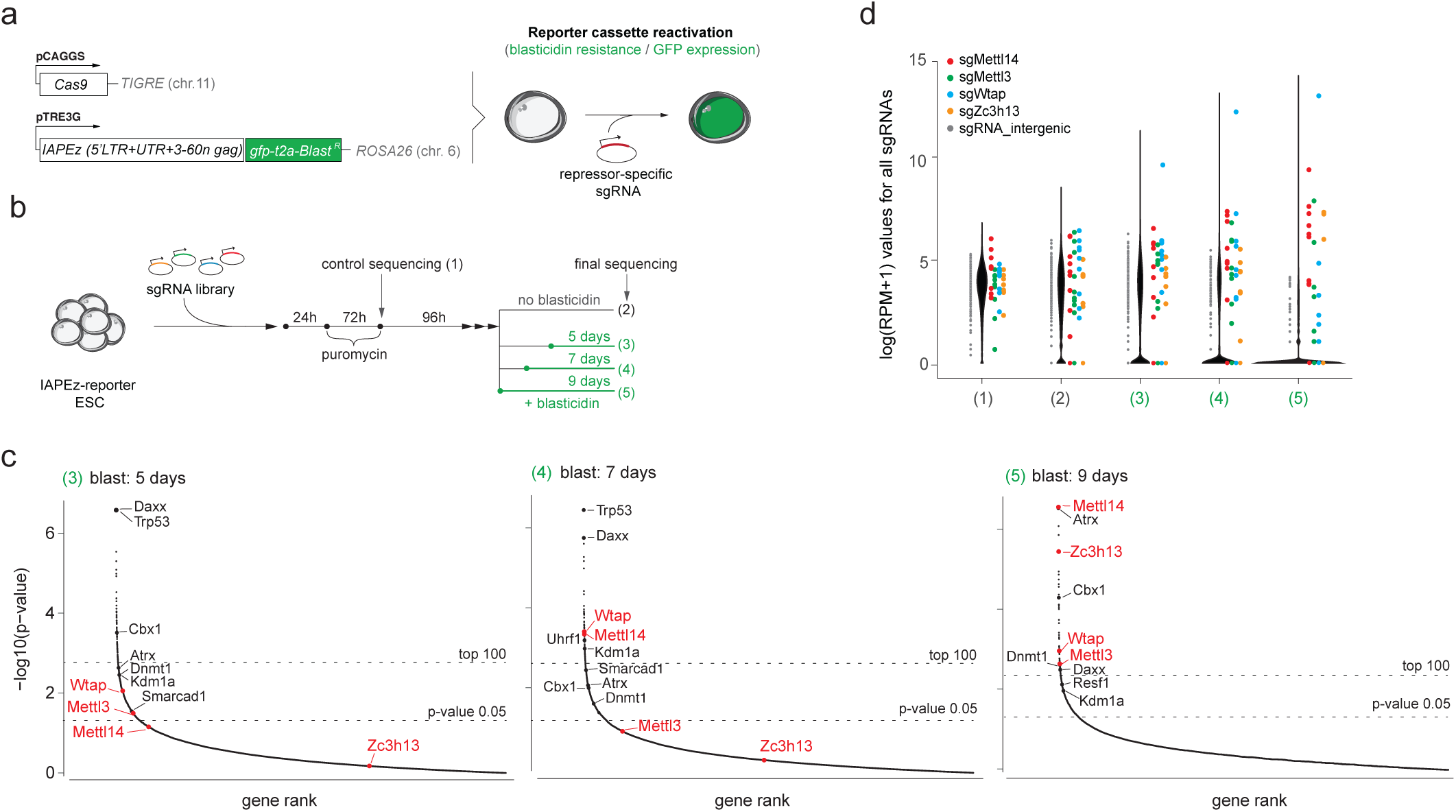
Genome-wide screen for IAP suppressors in mouse ESCs. **a**, Schematic of the IAPEz reporter cassette and experimental rationale. **b**, Workflow and timelines for the CRISPR-Cas9 screen. **c**, sgRNA ranking (MAGeCK) based on *P*-values for different selective pressures (5, 7 and 9 days of blasticidin treatment, as in (**b**). Dashed lines indicate the top 100 and *P*-value = 0.05 threshold. Known IAP regulators (black) and m^6^A methyltransferase complex members (red) are reported. **d**, Violin plots of sgRNA read counts, represented as the logarithm of Reads Per Million (RPM), upon library introduction (1), without selection (2) and upon blasticidin selection (3-5), as in (**b**); sgRNAs targeting *Mettl14* (red), *Mettl3* (green), *Wtap* (blue), *Zc3h13* (orange) and intergenic sequences (grey) are indicated.

For the screen, we transduced IAPEz-reporter cells with a lentiviral genome-wide sgRNA library at multiplicity of infection (MOI)= 0.2-0.3^12^ (Fig. 1b). Frequencies of sgRNAs upon blasticidin selection (5, 7 and 9 days) versus non-selected conditions were assessed via deep sequencing and candidate genes were identified using Model-based Analysis of Genome-wide CRISPR/Cas9 Knockout (MAGeCK)^13^. Efficiency of selection was evidenced by drop-out of control intergenic sgRNAs and genes were ranked based on sgRNA *P*-values (Fig. 1c, Supplementary Table 4, Supplementary Table 5). Although genome-wide screens of this magnitude typically suffer from low-statistical confidence, we identified several—but not all—genes previously associated with ERV repression (*Resf1, Trp53, Daxx, Atrx, Uhrf1, Cbx1, Dnmt1*)^14–17^ among the top 100 hits. Obtaining an incomplete list of known regulators is a common outcome of genome-wide screens which can be due to multiple factors including time-dependent dropout of essential genes with strong effects on cell viability (such as *Kap1*)^18^, heterogeneity in sgRNA efficiencies or limited representation of sgRNAs. Nonetheless, we were able to identify several novel candidates for IAP control (Supplementary Table 4), offering a foundation for future studies.

Notably, among the top hits were genes that encode regulators of the N^6^-Methyladenosine (m^6^A) mRNA methylation pathway, namely *Mettl3, Mettl14, Wtap* and *Zc3h13*, whose enrichment gradually increased with extended blasticidin-mediated pressure (Fig. 1c,d). By repeating the screen under the most stringent selection, we confirmed significant enrichment for *Mettl3, Mettl14* and *Wtap* sgRNAs (Extended Data Fig. 2a,b). m^6^A is the most abundant internal mark on eukaryotic mRNAs, critical for organizing mRNA fate— including export, decay and translation—in an array of biological processes such as embryonic development, cell differentiation, stress response and cancer^5^. Deposition of m^6^A is exerted co-transcriptionally by a nuclear multi-factor complex with an enzymatically active core formed by methyltransferase-like (METTL) METTL3 and METTL14 proteins and additional calibrating subunits, including Wilms tumor 1-associated protein (WTAP) and zinc finger CCCH-type containing 13 (ZC3H13). METLL3 and METTL14 form a heterodimer, wherein METTL3 is the catalytic component while METTL14 facilitates binding to the RNA substrate. WTAP and ZC3H13 are essential for the assembly of the complex into the nucleus^19^.

To confirm that the m^6^A RNA methylation pathway regulates endogenous IAP copies, we generated individual ESC lines of *Mettl3, Mettl14, Wtap* and *Zc3h13* gene knock-outs (KOs) (Fig. 2a, Extended Data Fig. 3a–d). To avoid the proliferation and differentiation effects that were previously reported in ESCs upon depletion of the mRNA methylation complex when cultured in metastable naïve conditions (serum+LIF), we derived and maintained our mutant lines in conditions that stabilize ground state pluripotency (2i+LIF) to preserve ESC identity^20^. When replacing ‘2i+LIF’ with ‘serum+LIF’ medium, we indeed observed drastic morphological changes in all KO lines (Extended Data Fig. 3e). Moreover, self-renewal ability was severely impaired in *Wtap-* or *Zc3h13*-KO cells upon serum conversion (Extended Data Fig. 3f). This could explain the low sgRNA ranking for these two genes in the screen upon extended selection time in ‘serum+LIF’ culture (Extended Data Fig. 2b).

**Fig. 2.**
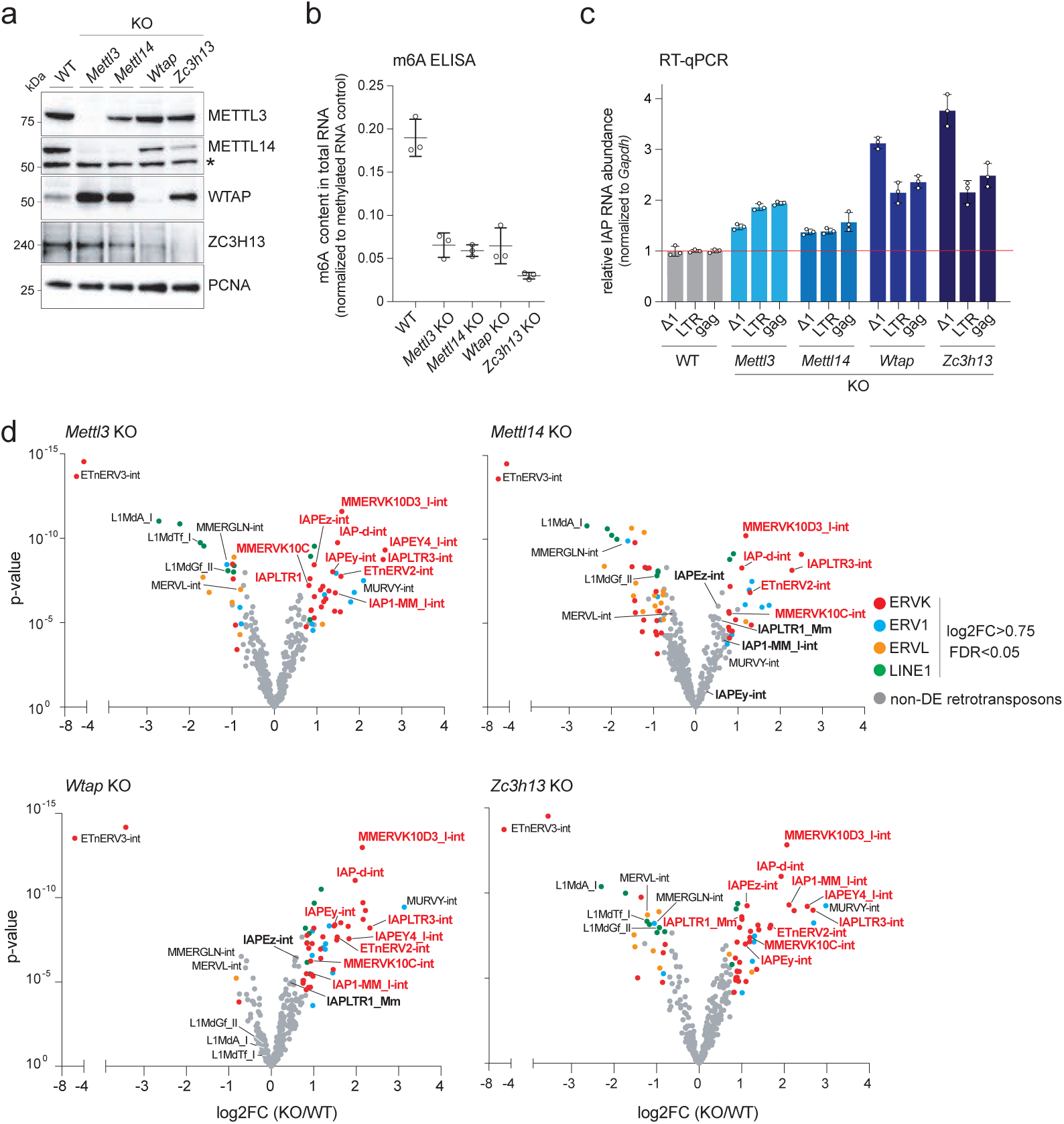
Depletion of the m^6^A methyltransferase complex increases IAP mRNA levels. **a**, Immunoblots showing *Mettl3*-, *Mettl14*-, *Wtap*- and *Zc3h13*-KO in ESCs. PCNA serves as a loading control. Asterisks indicates unspecific band **b**, Content of m^6^A in total RNA in WT and KOs as measured by m^6^A ELISA (data shown as means ± s.d. from three biological replicates). **c**, RT-qPCR analysis of steady-state IAP mRNA levels using primers against Δ1 elements and against LTR and *gag* sequences in WT, *Mettl3-, Mettl14-, Wtap-* and *Zc3h13-* KO (data shown as means ± s.d. from three biological replicates). IAP levels were normalized to *Gapdh* and expressed relative to WT levels (set to 1). **d**, Volcano plot representation of retrotransposon expression as measured by RNA-seq in *Mettl3-, Mettl14-, Wtap-* and *Zc3h13*-KO compared to WT. Red, blue, orange and green dots indicate significantly deregulated RepeatMasker annotations belonging to ERVK, ERV1, ERVL and L1 families, respectively (log2FC>0.75 and FDR<0.05). Non-differentially expressed (non-DE) retrotransposons are shown in grey. Names of upregulated ERVK elements are highlighted in red.

We first confirmed reduction in m^6^A RNA methylation levels in the four KO lines by ELISA (Fig. 2b). Then, we validated by RT-qPCR that knocking-out each of the four m^6^A factors increases endogenous IAPEz mRNA abundance by 2- to 3-fold compared to wild-type (WT) cells (Fig. 2c). RNA-seq analysis confirmed that IAPEz transcripts were significantly upregulated in the different m^6^A mutants, as well as close relatives within the ERVK family—namely MMERVK10C, MMERVK10D3, ETnERV2 and Y chromosome-specific IAPEy elements—that share above 65% of sequence identity with IAPEz (Dfam.org)(Fig. 2d, Extended Data Fig. 4a, Supplementary Table 6). Specific Y-linked elements of the ERV1 family were also more abundant in the KO lines (MuRVY). Notably, ERVL and long interspersed nuclear elements (LINEs or L1s) transcripts remained globally unaffected or downregulated (Fig. 2d and Extended Data Fig. 4b–d). These contrasted responses to the loss of m^6^A mRNA methylation highlight the divergent effects that this pathway may exert depending on the retrotransposon type, with a negative impact on IAP-related ERVK elements, specifically.

We then evaluated the potential impact of increased IAP transcripts on gene regulation. Upon reactivation, ERVs can affect the transcriptional state of neighboring genes through their LTR promoters and can modify the splicing pattern of host genes when located within introns. As previously documented, hundreds of gene transcripts were upregulated in m^6^A mutant ESCs^21^, among which 941 were common between the four KO lines (Extended data Fig. 5a). However, these upregulated genes did not show correlation with proximity of ERVK annotations, IAPs included (−5kb to +1kb from the Transcription Start Site, TSS) (Extended Data Fig. 5b). Moreover, we did not score increased occurrence of splicing between exonic sequences and IAP fragments in m^6^A mutants compared to WT ESCs (Extended Data Fig. 5c). As a whole, we conclude that the increased abundance of IAP transcripts upon m^6^A loss exhibits minimal *cis*-effects on gene expression. Importantly, depletion of m^6^A did not result in downregulation of known retrotransposon repressors and did not alter ESC identity, as demonstrated by the expression levels of pluripotency and early differentiation genes (Extended Data Fig. 5d,e). This provides strong indication that METTL3/14-dependent methylation directly represses IAP elements.

To prove this direct effect, we mapped the abundance and topology of m^6^A methylation on IAPEz transcripts, by performing m^6^A immunoprecipitation (MeRIP-seq)^22^ on total RNA from WT and *Mettl3-*KO ESCs in triplicates, with an average depth of 80M of sequenced fragments per sample. We scored 15,216 and 4,864 m^6^A peaks in WT and *Mettl3-*KO ESCs, respectively, with substantially higher m^6^A signal intensities in WT (Fig. 3a). A set of novel m^6^A peaks was detected in *Mettl3-*KO cells, however, they are likely false positive as they showed low signal intensity compared to canonical WT peaks (Fig. 3a, bottom right). Strikingly, in addition to the well-characterized enrichment of m^6^A methylation at the 3’UTR and exons of genic mRNAs^21–24^ (Fig. 3b and Extended Data Fig. 6a,b), we found that a significant number of METTL3-dependent m^6^A events mapped to retrotransposon annotations, comprising 13% of all peaks, including L1s—as previously reported^25,26^—and ERVK elements (Fig. 3b). When we plotted the distribution of m^6^A signal along the IAPEz consensus sequence specifically, we found two distinct regions of METTL3-dependent m^6^A enrichment, predominantly at the 5’UTR—present on the original IAP reporter—and to a lesser extent at the end of the *Pol* gene (Fig. 3c,d and Extended Data Fig. 6c). Enrichment in m^6^A also specifically coincided with the 5’UTR region of MMERVK10C elements (Extended Data Fig. 6d,e). Finally, m^6^A RNA methylation mostly occurs on conserved sequence motifs of RRACH (where R represents A or G, and H represents A, C or U)^22,23^. Accordingly, we found multiple RRACH motifs on the 5’UTR of IAPEz and MMERVK10C consensus sequences, half of them showing position conservation between the two element types (Extended data Fig. 6f). These data demonstrate for the first time that IAPs and their ERVK relatives undergo METTL3-dependent RNA methylation, uncovering a novel post-transcriptional pathway of retrotransposon suppression.

**Fig. 3.**
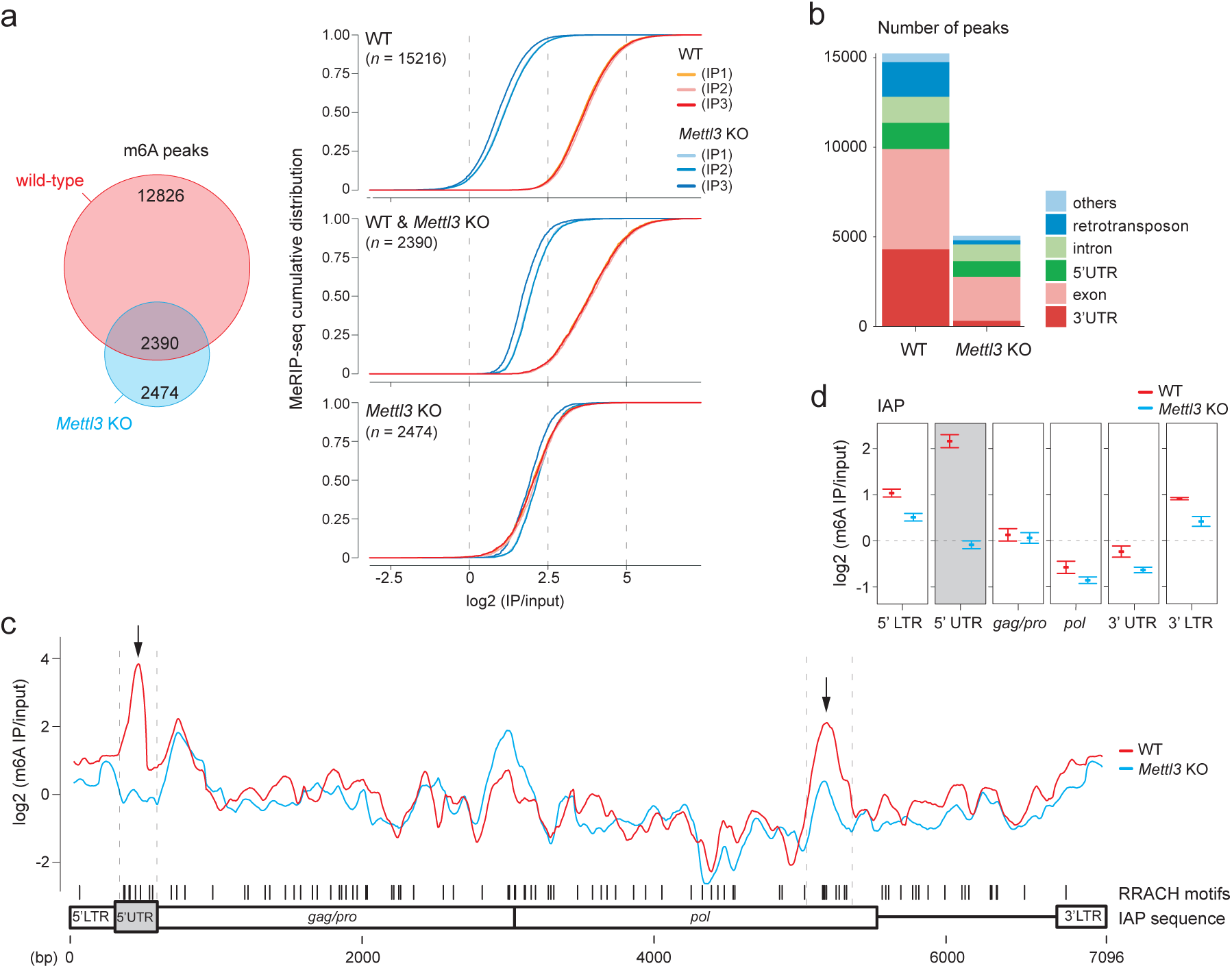
IAP mRNAs are directly targeted for m^6^A methylation. **a**, Left panel: Venn diagram shows overlap of m^6^A peaks between WT and *Mettl3-*KO ESCs identified by m^6^A RNA-IP; right panel: cumulative m^6^A signal intensity distribution (normalized to input) for m^6^A peaks in WT (top), overlapping (middle) and *Mettl3*-KO specific (bottom). **b**, Barplot demonstrating the number of peaks within indicated genomic features. **c**, Average of input-normalized m^6^A signal intensities for three m^6^A-IP replicates along the IAPEz consensus sequence in WT (red) and *Mettl3-*KO (blue). Distribution of RRACH motifs is indicated as vertical black lines. Black arrows point to regions of METTL3-dependent m^6^A enrichment (present in WT and lost in *Mettl3-KO*). **d**, Average of m^6^A signal intensities for the indicated IAP sequence segments (m^6^A-IP replicates, *n*=3).

Functions of the m^6^A RNA methylation complex have so far been investigated by conventional gene perturbation techniques -knock-out or shRNA-mediated knockdown-which precludes examining the early consequences of m^6^A loss and can additionally lead to secondary, confounding effects upon prolonged selection. To address the early and direct IAP responses to m^6^A depletion, we therefore used auxin-inducible degron (AID)^27^ to control acute and reversible depletion of the proteins forming the core of the m^6^A methyltransferase complex, METTL3 and METTL14, individually and in combination. Homozygous knock-ins were generated by introducing a 3xFLAG-AID cassette at the N-terminus of endogenous *Mettl3* and *Mettl14* loci in ESCs expressing the *Oryza sativa* TIR1 (*OsTir1*) E3 ligase, which mediates ubiquitination of AID-tagged proteins upon treatment with auxin (also known as indole-3-acetic acid, IAA) (Fig. 4a, Extended Data Fig. 7a,b). Addition of auxin prompted efficient and near-total degradation of AID-tagged METTL3 and METTL14 within 1 hour, which persisted over prolonged periods of auxin treatment and was readily reversible upon auxin washoff (Fig. 4b, Extended Data Fig. 4c). Depletion of METTLs was rapidly followed by significant and sustainable decrease of m^6^A RNA methylation levels (Fig. 4c). Most importantly, we observed a progressive, time-dependent accumulation of IAP transcripts upon m^6^A removal (Fig. 4d, Extended Data Fig. 7d). A similar trend was observed upon degron of the accessory m^6^A factor ZC3H13 (Extended Data Fig. 7e-h). After 96h of abolition of m^6^A-dependent regulation, IAP mRNA levels reached an 8-, 4- and 15-fold increase in single METTL3, single METTL14 or double METTL3; METTL14 degron, respectively (Fig. 4d), and this translated into accumulation of IAP-encoded GAG proteins in cytoplasmic speckles (Fig. 4e). Remarkably, IAP mRNA abundance was higher upon degron than in established single KO for individual *Mettl* genes (∼2-fold in *Mettl3-* or *Mettl14*-KO, Fig. 2c), suggesting the implementation of adaptive mechanisms upon prolonged m^6^A loss. Moreover, the relative upregulation in simultaneous compared to single degrons of METTL3 and METTL14 highlight their functional synergy in reducing IAP mRNA levels. Finally, re-stabilizing METTL proteins upon auxin removal prompted rapid decline in IAP mRNAs (Fig. 4d). Altogether, these results definitively argue that m^6^A RNA methylation dynamically restrains the cellular availability of IAP mRNA molecules.

**Fig. 4.**
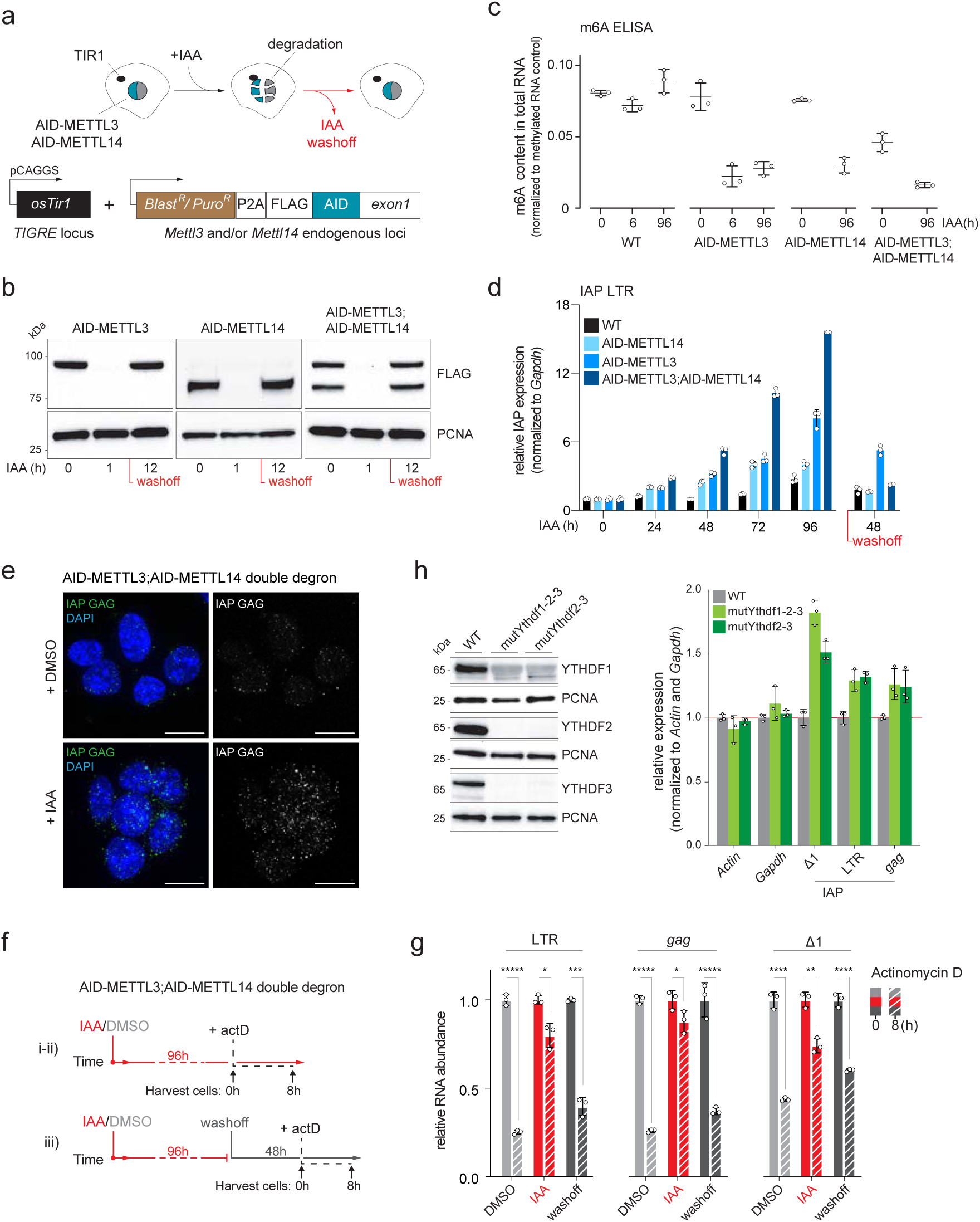
Auxin-inducible degron of METTL3 and METTL14 leads to increased IAP RNA levels and stability. **a**, Schematic of METTL3 and METTL14 degron engineering in ESCs. **b**, Immunoblot showing rapid and reversible auxin (IAA)-induced degradation of endogenous 3xFLAG-AID-tagged METTL3, METTL14 and METTL3; METTL4 (double degron, dd). PCNA serves as a loading control. **c**, ELISA showing m^6^A levels in total RNA after 0 and 96h of +IAA (treatment) of AID-METTL14, AID-METTL3 and AID-METTL3; AID-METTL14 dd (*n*=3 technical replicates). TIR1-only ESC line (hereafter donated TIR1) was used as WT control. **d**, RT-qPCR analysis of IAP mRNA levels using LTR-specific primers from 0 to 96h of IAA treatment, followed by 48h of IAA washoff in AID-METTL14 (light blue), AID-METTL3 (blue) and AID-METTL3; AID-METTL14 dd (dark blue). TIR1 line was used as WT control (data shown as means ± s.d. from three technical replicates). IAP levels were normalized to *Gapdh* and expressed relative to levels at 0h (set to 1). **e**, Immunofluorescence staining for IAP-GAG protein in AID-METTL3; AID-METTL14 dd without (left panel) and upon 96h IAA (right panel) treatment. **f**, Schematic of evaluation of IAP mRNA degradation rate in AID-METTL3; AID-METTL14 dd treated with either DMSO (control) or IAA for 96h and actinomycin D (actD) for 8h prior to (i-ii) or after 48h IAA/DMSO washoff (iii). **g**, RT-qPCR analysis of IAP mRNA levels in DMSO+actD (light grey), IAA+actD (red) and washoff+actD (dark grey) using LTR-, *gag*-, or Δ1-specific primers (data shown as means ± s.d. from three biological replicates, two-sided Student’s t-Test, asterisks indicate *P*-value significance). Simultaneous degradation of METTL3 and METTL14 leads to reversible increase in IAP mRNA stability. **h**, (Left) Immunoblots of mutYthdf1-2-3 and mutYthfd2-3 ESCs (see **Extended Data Fig. 8**). PCNA serves as a loading control. (Right) RT-qPCR shows increased IAP mRNA levels in mutYthdf1-2-3 and mutYthfd2-3 compared to WT using Δ1-, LTR- or *gag*-specific primers (data shown as means ± s.d. from three biological replicates). IAP levels were normalized to the geometric mean between *Actin* and *Gapdh* and relative to WT levels (set to 1). *Actin* and *Gapdh* were normalized to each other.

As m^6^A RNA methylation plays an essential role in tuning mRNA lifetime^28^, we considered that m^6^A could destabilize IAP-derived mRNAs. To test this, we measured IAP mRNA levels after blocking transcription for 8h with actinomycin D, in absence or in presence of the METTL core complex (96h of auxin versus DMSO treatment of double AID-METTL3; AID-METTL14 cells) (Fig. 4f). IAP mRNA levels were significantly higher in auxin-treated versus DMSO control (Fig. 4g), indicating that IAP mRNAs have an extended half-life in absence of the METLL complex. Importantly, the effect was reversible, as METTL restoration upon auxin removal decreased IAP mRNA levels (Fig. 4g, dark grey bars). The destabilization and decay of m^6^A-modified mRNAs is determined by a set of readers, among which the YTH-domain containing proteins YTHDF1, YTHDF2 and YTHDF3 play key role by directing modified mRNAs to dedicated cytosolic compartments^6^. We therefore reasoned that, similarly to depleting the m^6^A writing complex, depleting the YTHDF m^6^A readers should increase IAP mRNA abundance. To account for potential redundancy between YTHDF proteins, we aimed to simultaneously knocking-out the three genes by CRISPR-Cas9 targeting. We were able to acquire two mutant ESC lines that exhibited complete loss of all three proteins (Fig. 4h, left and Extended Data Fig. 8). Importantly, IAP mRNA levels were increased in both *Ythdf* mutant lines (Fig. 4h, right), underscoring that the m^6^A methylation pathway is involved in IAP mRNA clearance.

Regulation of ERV mRNA fate by m^6^A modification may be particularly relevant in situations of relaxation of chromatin-based control, such as happening in early mammalian embryos-from which ESCs are derived- or ageing. The link we uncovered with YTHDF m^6^A readers indicates that ERV mRNA decay occurs through YTHDF-promoted phase-partitioning into cytoplasmic processing bodies (P-bodies)^29^ and stress granules, which is congruent with previous observations that IAP mRNAs localize to these structures^30^. Additionally, this pathway may cooperatively prevent m^6^A-modified ERV mRNAs from being reverse transcribed, translated or assembled into retrotransposition complexes, providing multiple layers of control. To wit, we found that m^6^A mostly occurs on the 5’UTR of IAPs and their relatives, a region that contains the tRNA primer binding site (PBS) essential for reverse transcription. Finally, another role of m^6^A-dependent regulation could be to dampen the immunogenic potential of ERV-derived RNA species and their ability to trigger inflammatory responses, as seen in several neurodegenerative diseases linked to human ERV reactivation.

## Methods

### Cell culture

E14 mouse embryonic stem cells (ESCs) were grown in two different media: ‘serum+LIF’ consisted in Glasgow medium (Sigma), 15% FBS (Gibco), 2mM L-Glutamine, 0.1mM MEM non-essential amino acids (Gibco), 1mM sodium pyruvate (Gibco), 0.1mM b-mercaptoethanol, 1000 U/mL leukemia inhibitory factor (LIF, Miltenyi Biotec); ‘2i+LIF’ was made of 50% Neurobasal medium (Gibco), 50% DMEM/F12 (Gibco), 2mM L-glutamine (Gibco), 0.1mM β-mercaptoethanol, Ndiff Neuro-2 medium supplement (Millipore), B-27 medium supplement (Gibco), 1000 U/mL LIF, 3μM Gsk3 inhibitor (CT-99021), 1μM MEK inhibitor (PD0325901). Cells were cultured in 0.2% gelatin-coated flasks at 37 °C with 5% CO_2_. Except for the CRISPR-Cas9 loss-of-function screens that were performed in serum-based medium, all experiments were performed in 2i medium.

### Plasmid construction

#### IAPEz reporter

Expression vectors targeting *ROSA26* (pEN111) and *TIGRE* (Addene plasmid #92141) loci and the *ROSA26*- and *TIGRE*-specific sgRNA encoding plasmids were kindly provided by Elphége Nora (UCSF). The IAPEz-5’LTR-5’UTR-gag(3-60nt) consensus sequence was obtained from RepeatMasker and/or from repbase (http://www.repeatmasker.org/) synthetized and cloned into pUC57 by GenScript (Supplementary Table 1). To make the IAPEz reporter (pTCH1), we combined IAPEz-5’LTR-5’UTR-gag(3-60nt) fragment and the GFP-T2A-blasticidin cassette using extension PCR and inserted it into the pEN111 backbone using *ClaI* site. To insert Cas9 gene at the *TIGRE* locus, 3xFLAG-NLS-Cas9 was amplified from pX459 expression vector (Addgene plasmid #62988) and inserted into #92141 backbone using *BamHI* and *XhoI* sites (pTCH2).

#### Plasmids for N-terminus tagging with AID domain for auxin-inducible degron

Vectors for targeting AID inserts into gene N-terminus were generated as follows: either PuroR-P2A-3xFLAG-AID, or BlastR-P2A-3xFLAG-AID inserts were cloned into pUC19 backbone (pFD71 with puromycin resistance gene and pFD75 with blasticidin S resistance gene). Next, homology arms ranging from 320pb to 530bp depending on the gene (flanking both sides, but excluding ATG start codon) for *Mettl3, Mettl14* and *Zc3h13* were PCR amplified from mouse genomic DNA and inserted into pFD71 or pFD75 surrounding and in frame with AID insert using *EcoRI/NcoI* sites for upstream homology arms and *AgeI/HindII* for downstream homology arms. Final expression vectors were used as follows: pTCH3 (BlastR-P2A-3xFLAG-AID-METTL3_Nter), pTCH4 (BlastR-P2A-3xFLAG-AID-METTL4_Nter) and pFD119 (BlastR-P2A-3xFLAG-AID-ZC3H13_Nter) were used to generate individual endogenous degron lines for aforementioned genes, pTCH4 and pTCH5 (PuroR-P2A-3xFLAG-AID-Mettl3) were used sequentially to generate METTL3;METTL14 double degron cell line. For sgRNA cloning, the pX459 plasmid (Addgene #62988) was digested with *BbsI* immediately downstream of the U6 promoter and annealed DNA duplex corresponding to the target sgRNA sequences were ligated. sgRNA sequences were chosen to overlap with gene TSS, so that upon AID insert introduction the sgRNA-specific sequences were disrupted. sgRNA sequences used for degron targeting are listed in Supplementary Table 2.

#### Cell transfection, clone isolation cell line validation

All transgenic insertions and mutations were performed using Amaxa 4D nucleofector (Lonza). For each nucleofection, 3-5 × 10^6^ cells were electroporated with 1-3ug of non-linearized targeting vector and/or sgRNA/Cas9-encoding plasmids and plated at a low density. Two days later, cells were selected with puromycin (1μg/mL, Life Technologies) or blasticidin S (5μg/mL) for 2 or 5 days (for generation of knock-out and knock-in cell lines respectively) and individual clones were picked and screened by PCR. Flippase-mediated removal of puromycin resistance cassettes were performed for the IAPEz reporter cell line (from both *ROSA26* and *TIGRE* loci) and for puromycin resistance cassette for METTL3;METTL14 double-degron from *TIGRE* locus. For the IAPEz reporter cell line, functionality of the reporter cassette was confirmed by dox-induced expression followed by FACS and fluorescence microscopy while Cas9 expression and activity was confirmed by *Kap1*-specific sgRNA introduction (see below) and western blot analysis. To generate *Mettl3, Mettl14, Wtap, Zc3h13-* KO and *Ythdf* mutant ESCs, two sgRNAs for each gene were designed using the online CRISPOR Design Tool^31^ to introduce deletions. For sgRNA cloning, the pX459 plasmid (Addgene #62988) was digested with *BbsI* immediately downstream of the U6 promoter and annealed DNA duplex corresponding to the target sgRNA sequences were ligated. The *Mettl3*-KO cells were created by deleting part of exon 4; *Mettl14*-KO by deleting part of exon 1; *Zc3h13*-KO by deleting part of exon 9; and *Wtap*-KO by deleting exon 3 and part of exon4. *mutDF1-2-3* and *mutDF2-3* ESCs were created by simultaneous introduction of six sgRNAs targeting *Ythdf1, Ythdf2* and *Ythdf3* genes. For *Mettl3-, Mettl14-, Zc3h13*- and *Wtap*-KO, deletions were confirmed by Sanger sequencing, western blot analyses and m^6^A ELISA. For *mutDF1-2-3* and *mutDF2-3* mutations were confirmed by Sanger sequencing and western blot analyses. For degron lines, proper insertion and AID-fusion protein activities were confirmed by genotyping, western blot and m^6^A ELISA. For sgRNA sequences used for generation of KO, mutant and degron lines see Supplementary Table 2.

#### Lentivirus production and lentiviral-based *Kap1*-specific sgRNA knockout

Two previously described sgRNAs specific to the *Kap1* gene^10^ (Supplementary Table 2) were incorporated into plentiGuide-puro vector (Addgene #52963). For production of lentiviral particles, HEK293FT cells were co-transfected with 3.33µg of either of the sgKap1 lentiGuide-puro constructs, 2.5µg psPAX2 packaging plasmid and 1µg pMD2.G envelope plasmid using Lipofectamine 2000 (Invitrogen). Lentiviral supernatant was collected, filtered with 0.45μm filter, concentrated using Amicon Ultra centrifugal filter (Millipore, 100kDa cutoff) and added to pre-plated IAPEz reporter cells supplemented with 8µg/ml polybrene (Millipore). Twelve hours after infection the medium was replaced and supplemented with puromycin (1μg/mL). After 48 hours of puromycin selection the media was replaced and supplemented with blasticidin S (5μg/mL) for additional 72 hours.

#### Western blotting

Cells were lysed for 20 min on ice using RIPA buffer (1xPBS, 0.5% sodium deoxycholate, 0.1% sodium dodecyl sulfate, 1% Igepal) supplemented with protease inhibitors. Cellular debris were spun down for 20 min at 16,400 rpm, at 4°C and supernatant was collected. Protein concentration was assessed using Bradford assay. Next, protein extracts were denaturated in LDS loading buffer (Life Technologies) supplemented with 200mM DTT and boiled for 10 min at 95°C. Equal amounts of protein extracts were loaded on 4-12% Bis-Tris gel (NuPAGE) or 3-8% Tris-Acetate gel (NuPAGE) for ZC3H13 detection. Proteins were transferred on 0.45μm nitrocellulose membrane (GE Healthcare), blocked with 5% milk with PBS-Igepal (PBS + 0,3% Igepal) for 1 hour at room temperature (RT). Membranes were incubated with primary antibody (Supplementary Table 3) at 4°C overnight in 1% milk, washed 5 times with 0.3% PBS-Igepal and incubated with HRP-conjugated secondary antibodies for 1 hour at RT and washed again 5 times with 0.3% PBS-Igepal. Signal was detected using LumiLight Plus Kit (Roche) on the Chemidoc MP imaging system (Biorad).

#### FACS analysis

Cells were collected, washed with PBS to remove residual medium and proceeded to analyze GFP expression using NovoCyte Flow cytometer (ACEA Biosciences). The percentage of GFP-positive cells was determined upon definition of three gates: i) FSC-H vs SSC-H to isolate cells from debris,ii) SSC-H vs SSC-A to isolate single cells and iii) SSC-H vs FITC-H for detection of GFP-positive population.

#### Genome-wide screen in IAPEz reporter mouse ES cells (Screens I and II)

Approximately 300 × 10^6^ IAPEz reporter cells expressing Cas9 were lentivirally infected with a genome-wide Mouse Two Plasmid Activity-Optimized CRISPR Knockout Library (Addgene #1000000096) as described above, containing 188,509 sgRNAs targeting 18,986 genes and 199 intergenic sgRNAs at a multiplicity of infection of 0.2–0.3 (measured by puromycin resistance gene co-delivered with the lentiviral vector) and selected for lentiviral integration using puromycin (1μg/mL) for 3 days. In Screen I, the culture was expanded for another 4-6-8 days. On day 4, 6 and 8 of expansion, 200 × 10^6^ cells were split into blasticidin S selecting conditions (for 9, 7 and 5 days respectively) and non-selection conditions (9 days). Cells in non-selection conditions were maintained at minimum level of 100 × 10^6^ cells and logarithmic growth. After 9 days 3-5 × 10^6^ cells from selection conditions and 100 × 10^6^ non-selection conditions were washed 3 times with PBS and pelleted by centrifugation for genomic DNA extraction using GeneElute Mammalian Genomic DNA Miniprep kit (Sigma) and Quick-DNA Midiprep Plus kit (Zymo Research) respectively following the manufacturers guidelines. The sgRNA-encoding insertions were PCR-amplified using Agilent Herculase II Fusion DNA Polymerase (600675). These libraries were then sequenced Illumina HiSeq (approximately 5-10 million reads with sgRNA sequence per condition; around 40x coverage per library element in non-selection conditions, Screen I). Based on the results from Screen I, demonstrating that the longer blasticidin S treatment, the better intergenic sgRNA depletion, we performed Screen II in two biological replicates with 9-day long blasticidin S selection following either 8-day long, or 17-day long cell culture (<u>early</u> and <u>late</u> selection respectively). Upon genomic DNA extraction and library amplification libraries were sequenced using an Illumina HiSeq 2500 (SE65) (approximately 30-35 million reads per condition; around 170x coverage per library element in early and late non-selection conditions, Screen II. See Supplementary Table 1 for the primer sequences used to amplify the libraries.

#### Immunofluorescence

Cells were plated on fibronectin-coated (Sigma) glass cover slips. For IAPEz reporter reactivation control, the medium was supplemented with doxycycline (1µg/mL) for 24 hours. The next day, cells were fixed with 3% paraformaldehyde for 10 min at room temperature, rinsed three times with PBS, incubated 3 min in 0.3 μg/mL DAPI and rinsed again with PBS. For detection of IAP GAG, after fixation, cells permeabilized for 4 min with PBS/0.5x-Triton-X100 on ice, blocked with 1% BSA/PBS for 15 min, incubated for 40 min with rabbit anti-mouse IAP-GAG antibody (Gift from B. Cullen), 40 min with secondary antibodies and 3 min in 0.3 μg/mL DAPI at RT. Slides were mounted with VECTASHIELD mounting media (Vector Laboratories). Images were obtained with an Upright Spinning disk Confocal Microscope (Roper/Zeiss).

#### RT-qPCR analysis

Total RNA was extracted using Trizol (Life Technologies). Genomic DNA was removed by DNase I treatment (Qiagen), precipitated and resuspended in DNase/RNase-free water. Next, 10µg of RNA was used for a second round of purification using RNeazy Mini columns (Qiagen) and 500ng RNA was reverse transcribed using random priming with Superscript III (Life Technologies). Real-time quantitative PCR was performed using the SYBR Green Master Mix on the Viia7 thermal cycling system (Applied Biosystem). Relative expression levels were normalized to the *Gapdh* using the ΔΔCt method. Error bars indicate standard deviation of 3 biological replicates for all the knock-out lines and 3 technical replicates for the degron cell lines. For primer sequences, see Supplementary Table 1.

#### Auxin (IAA) treatment

The *osTir1* (wild-type), METTL3, METTL14, ZC3H1313 single degron and METTL3;METTL14 double degron ESCs were plated on gelatin-coated 6-well plates (0.3× 10^6^ cells/well) or 100mm Petri dishes (1× 10^6^ cell/plate) depending on the experiment. On the next day, medium was replaced to fresh medium with 500µM IAA for 1-6-24-48-72-96 hours depending on the experiment. Fresh medium supplemented with IAA was replaced every 24 hours and cells were passaged every 48 hours. For the 48-hour washoff conditions the cells were washed with PBS and the medium was replaced with fresh medium without IAA. After 24 hours of recovery, cells were passaged and plated for expansion for another 24 hours. Next, total protein or RNA extracts were obtained (see above).

#### RNA stability assay

For RNA stability assay, 0.5 × 10^6^ of METTL3;METTL14 double degron ESCs treated with either IAA, DMSO or IAA washoff, were plated on fibronectin-coated 6cm plates. 24 hours later, the medium was replaced to fresh medium supplemented with 5µM actinomycin D (Sigma) to inhibit transcription. Total RNA was extracted at indicated time points and used for RT-qPCR.

#### Poly-A RNA Sequencing

Total RNA was extracted using Trizol (Life Technologies). Genomic DNA was removed by in solution DNaseI treatment (Qiagen), RNA was precipitated and resuspended in DNase/RNase-free water. Next, 10µg of RNA was used for a second round of purification using RNeazy Mini columns (Qiagen). RNA integrity was evaluated on TapeStation (Agilent) using RNA ScreenTape (5067-5576,), requiring a minimal integrity number (RIN) of 9. Libraries were prepared according to Illumina’s instructions accompanying the *TruSeq Stranded mRNA Library Prep Kit* (Illumina). 800ng of RNA per replicate was used for library preparation. After library preparation, the length profiles were assessed with the LabChip GX Touch HT system (Perkin Elmer) and equimolar pool from all samples was prepared. Molarity of the pool was quantified by qPCR using KAPA Library Quantification Kit and the CFX96 qPCR system (Biorad) before sequencing. Samples were sequenced using Novaseq 6000 (PE100, approx. 80M clusters per replicate).

#### m^6^A IP and Sequencing

m^6^A IP was carried out using Magna MeRIP m^6^A Kit (Millipore) according to the manufacturer’s instructions. In short, total RNA was extracted using Trizol (Life Technologies). DNaseI-treated and RNA samples were chemically fragmented into 100-nucleotide-long fragments and 350µg of total RNA were subjected to each immunoprecipitation (IP) with affinity purified anti-m6A antibody in presence of RNase inhibitor. Bound m6A-methylated RNA fragments were eluted with free N6-methyladenosine, purified using RNeazy Kit (Qiagen) and processed for library generation using *SMARTer Stranded Total RNA-Seq Kit v2 - Pico Input Mammalian* (TaKaRa) following the manufacturer’s recommendations, but without fragmentation step (9ng of RNA per replicate). Sequencing was performed using Illumina Novaseq 6000 System. The m^6^A IP for WT and *Mettl3-*KO cells were performed in three technical replicates. Input for each of the cell lines was sequenced as a control.

#### Genome-wide screen analysis

The sequenced reads were mapped to the sgRNA library. Only reads that contained one sgRNA sequence without mismatch were counted. The MAGeCK^13^ test command line (version 0.5.8) was used to rank sgRNAs and genes with following parameters: *--norm-method total --adjust-method fdr --remove-zero-threshold 10 --gene-lfc-method alphamean -- remove-zero both*.

#### RNA-seq analysis

Adapters were trimmed using Atropos v1.1.16^32^. The paired-end reads alignment was performed onto the Mouse reference genome (mm10) with STAR v2.7.0a^33^ reporting randomly one position, allowing 6% of mismatches (*--outFilterMultimapNmax 5000 -- outSAMmultNmax 1 --outFilterMismatchNmax 999 --outFilterMismatchNoverLmax 0.06*). Repeats annotation was downloaded from RepeatMasker (http://www.repeatmasker.org/). In order to reconstruct full-length LTR copies, we used the same strategy as done previously^10^ using the perl tool “*One code to find them all*”^34^. Reconstructed transposons annotation and basic genes annotation from GENCODE v18 were merged and used as input for quantification with featureCounts v1.5.1^35^. Differential expression analysis was performed using edgeR’s normalisation combined with voom transformation from limma R package^36,37^. *P*-values were computed using limma and adjusted with the Benjamini-Hochberg correction. Genes and transposon families were declared as differentially expressed if FDR<5% and log2 Fold-change > 0.75. Upregulated genes in all four KO lines were annotated with proximal retrotransposon elements (overlap with promoter regions defined as -5kb to +1kb from the TSS). Randomized gene sets were created 100 times and were annotated to proximal retrotransposon elements in order to compute permutation test using regioneR^38^ R package.

To estimate intron retention between genes and single IAP copies, reads alignment was performed using specific parameters to report only uniquely mapped reads with STAR v2.7.0a^33^ (*--outFilterMultimapNmax 1 --outSAMmultNmax 1*). Unannotated splice junctions detected by STAR was annotated with GENCODE v18 and IAP LTR elements from RepeatMasker annotation to retrieve splicing events between a gene and an IAP transposon. The number of uniquely mapped reads crossing the splicing events was calculated for each sample and normalized by the library size.

#### MeRIP-seq analysis

Due to the addition of 3 nucleotides on 5’end of the second sequencing read (R2) from the Pico v2 SMART Adapter, paired-end reads were trimmed using Trim Galore v0.4.4 with the options: *--three_prime_clip_R1 3 --clip_R2 3* (http://www.bioinformatics.babraham.ac.uk/projects/trim_galore/) Reads were aligned onto the mouse ribosomal sequence (GenBank: BK000964.3) using Bowtie v1.2 allowing at most 3 mismatches^39^. Previously unmapped reads were aligned onto the mouse reference genome (mm10) using STAR v2.6.0c reporting randomly one position, allowing 4% of mismatches (*--outFilterMultimapNmax 5000 --outSAMmultNmax 1 --outFilterMismatchNmax 999 --outFilterMismatchNoverLmax 0.04*). PCR duplicates were removed using STAR with the option *--bamRemoveDuplicatesType UniqueIdenticalNotMulti*. Bigwig files were produced with deepTools v2.5.3^40^ using the option *--normalizeUsingRPKM*. Peaks enriched in the MeRIP sample over the input control were defined using MACS2 peak-caller^41^ with a genome size of 994,080,837bp^21^ and the FDR threshold of 5%. Reads were extended to 200bp-long fragments. Only peaks called in at least two replicates were used for downstream analysis. Peaks intensity was calculated using featureCounts v1.5.1 and normalized to the background (reads not falling into peaks) and to the peak length. GENCODE v18 was used to define 5’UTR, 3’UTR, intronic and exonic regions. Retrotransposons annotation (RepeatMasker) was downloaded from UCSC table browser. Genes overlapping with at least one peak were used to calculate coverage along the genic region (5’UTR, coding sequence and 3’UTR) with trumpet R package^42^. Mapped reads onto the Mouse reference genome overlapping with IAP elements were extracted as single-end reads and mapped to the full-length IAP consensus sequence (GenBank: M17551.1) with Bowtie2 v2.2.9^43^ with these parameters: *--local -N 1*. Coverage along the consensus sequence was normalized to background (reads not falling into peaks) as it was done previously for peak intensities. Rolling mean was calculated for a window of 50bp in order to smooth the signal. RRACH motif was searched into the IAP consensus sequence using RSAT dna pattern.

## Extended Data Figure Legends

**Extended Data Fig. 1.**
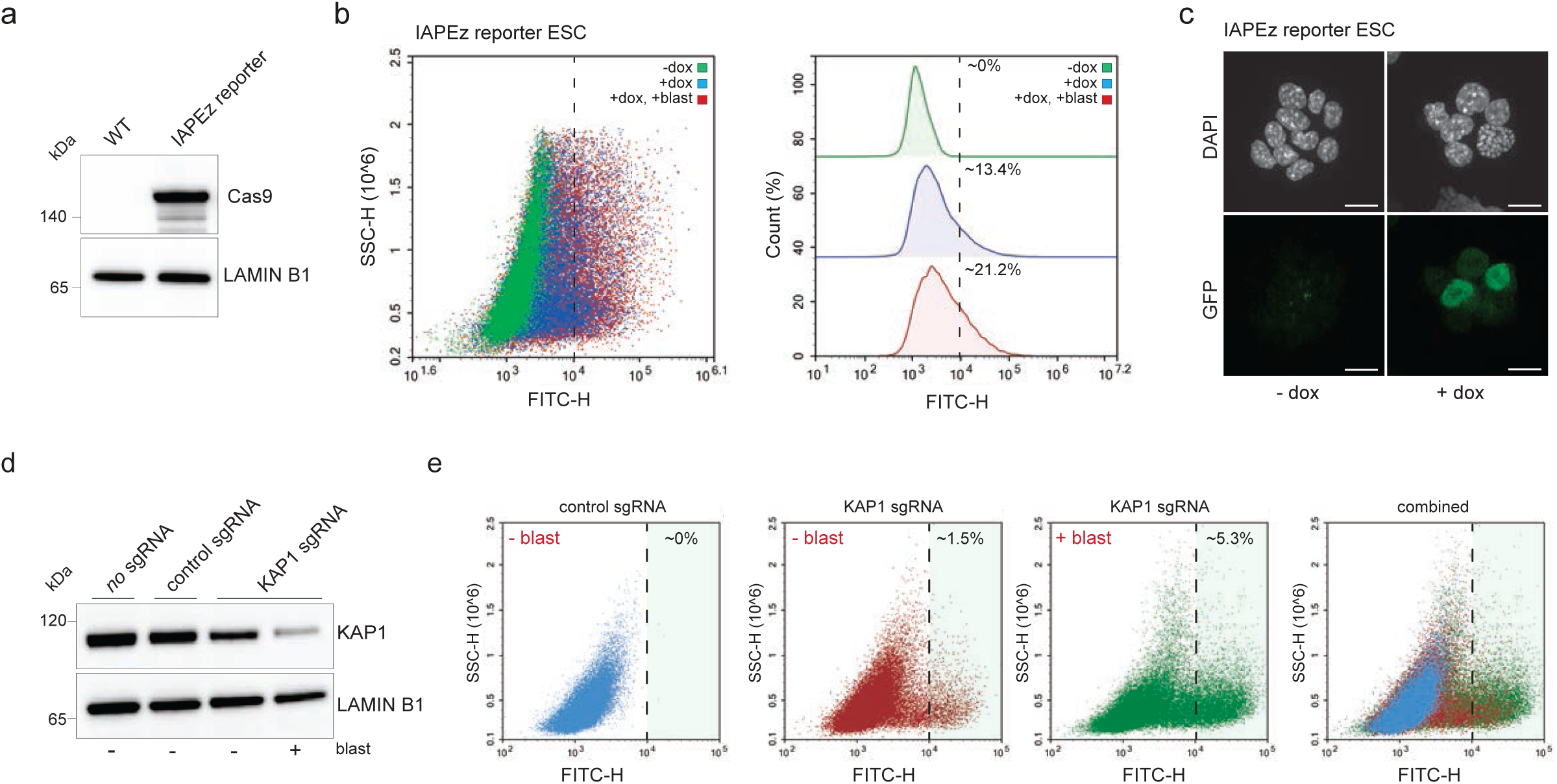
IAPEz reporter cell line validation. **a**, Immunoblot showing Cas9 levels in the parental (E14) and dox-inducible IAPEz reporter ESCs. LAMIN B1 serves as a loading control. **b**, Flow cytometry analysis of IAPEz reporter cassette activation upon dox induction and blasticidin selection. Analysis was performed using NovoExpress software (Acea Biosciences) **c**, Dox-induced IAPEz reporter activity analysis by fluorescence microscopy. **d**, Immunoblot showing KAP1 levels in IAPEz reporter ESCs upon KAP1-specific sgRNA introduction and blasticidin selection. Scrambled sgRNA was used as a control. Blasticidin treatment shows that only cells with successful KAP1 depletion become antibiotic-resistant. LAMIN B1 serves as a loading control. **e**, Flow cytometry analysis for GFP expression in cells from (**d**). KAP1 depletion combined with blasticidin selection leads to increase in GFP-positive cells.

**Extended Data Fig. 2.**
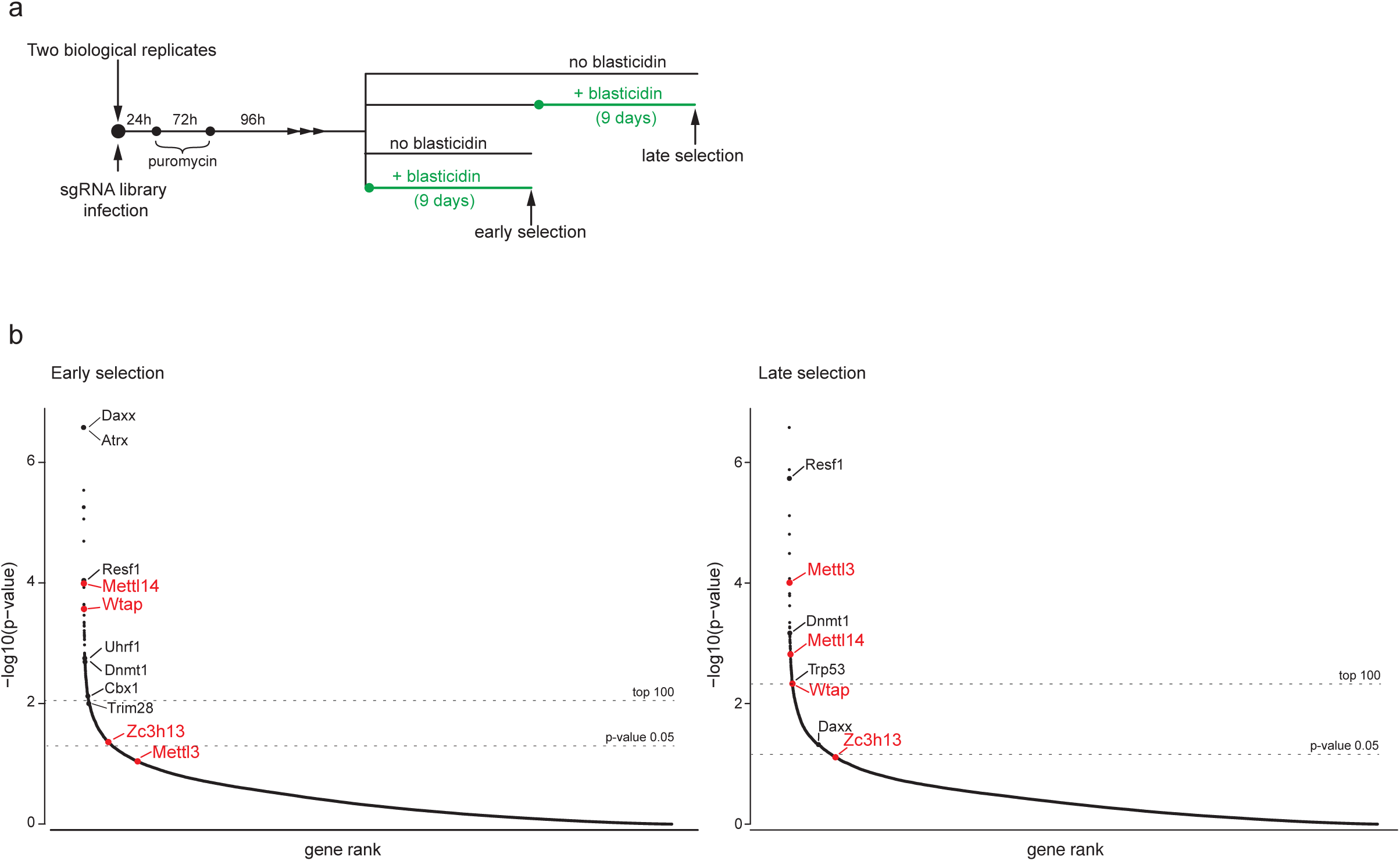
Early vs late genome-wide screen for IAP suppressors. **a**, Schematic of the screening process. The screens have been performed in two biological replicates. **b**, sgRNA ranking (MAGeCK) based on *P*-values for early (left) and late (right) blasticidin selection. Dashed lines show the top 100 and *P*-value=0.05 thresholds. Previously identified IAP regulators (black) and m^6^A methyltransferase complex members (red) are shown.

**Extended Data Fig. 3.**
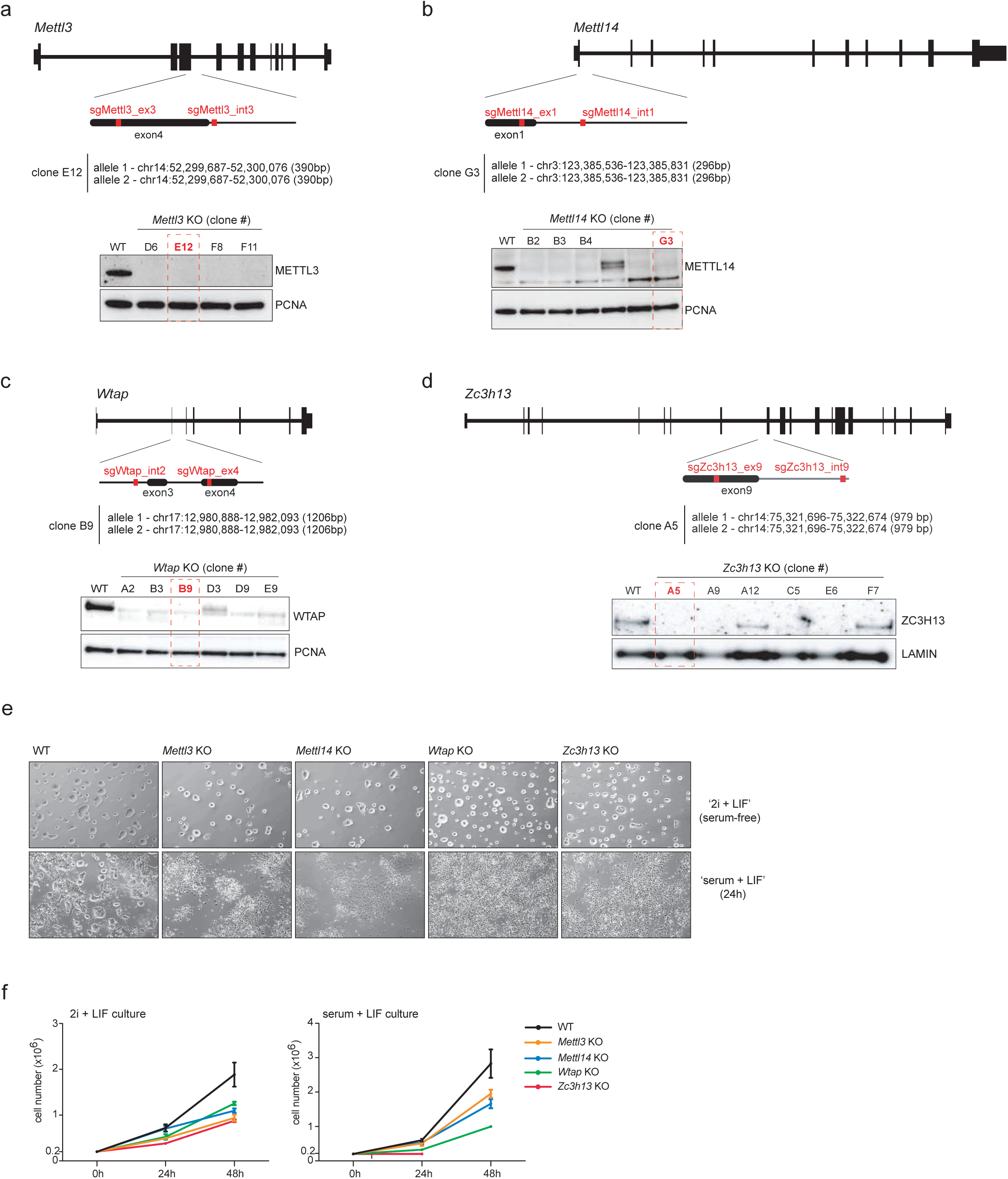
Generation of *Mettl3, Mettl14, Wtap* and *Zc3h13* knock-outs in mouse ESCs. **a-d**, *Mettl3-* (**a**), *Mettl14-* (**b**), *Wtap*- (**c**), and *Zc3h13*-KO (**d**), generated by CRISPR-Cas9 deletions. Schematic representation showing sgRNA sequences used to target indicated loci (top). Deletion for each locus was validated by genotyping, Sanger sequencing and western blot analyses (bottom, PCNA and LAMIN B1 serve as a loading controls) confirming ablation of indicated protein. In red, clones selected for downstream experiments. **e**, Bright field microscopy of indicated KOs grown in serum-free ‘2i+LIF’ medium, or upon conversion to ‘serum+LIF’ medium. **f**, Growth rate curves of WT and KOs. Note that for *Zc3h13*-KO cells are not stable in ‘serum+LIF’ medium.

**Extended Data Fig. 4.**
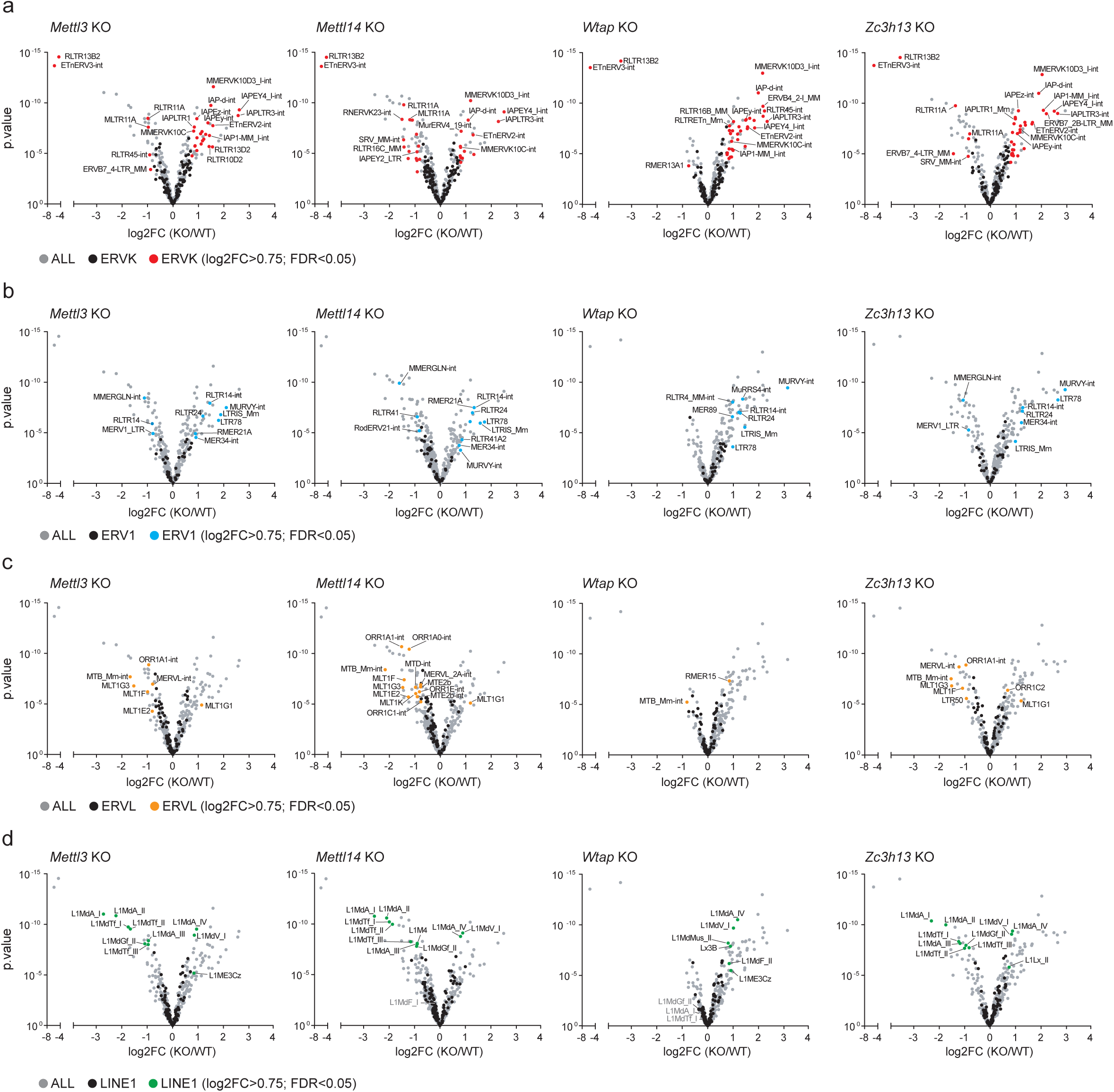
Depletion of m^6^A methyltransferase complex results in deregulation of different retrotransposon families. **a-d**, Volcano plot representations of up- and down-regulated ERVK (**a**), ERV1 (**b**), ERVL (**c**) and LINE-1 (**d**) retrotransposons measured by RNA-seq in *Mettl3-, Mettl14-, Wtap- and Zc3h13-*KO compared to WT. Red, blue, orange and green dots indicate significantly deregulated repeats for indicated families (log2FC>0.75 and FDR<0.05). Annotations from RepeatMasker.

**Extended Data Fig. 5.**
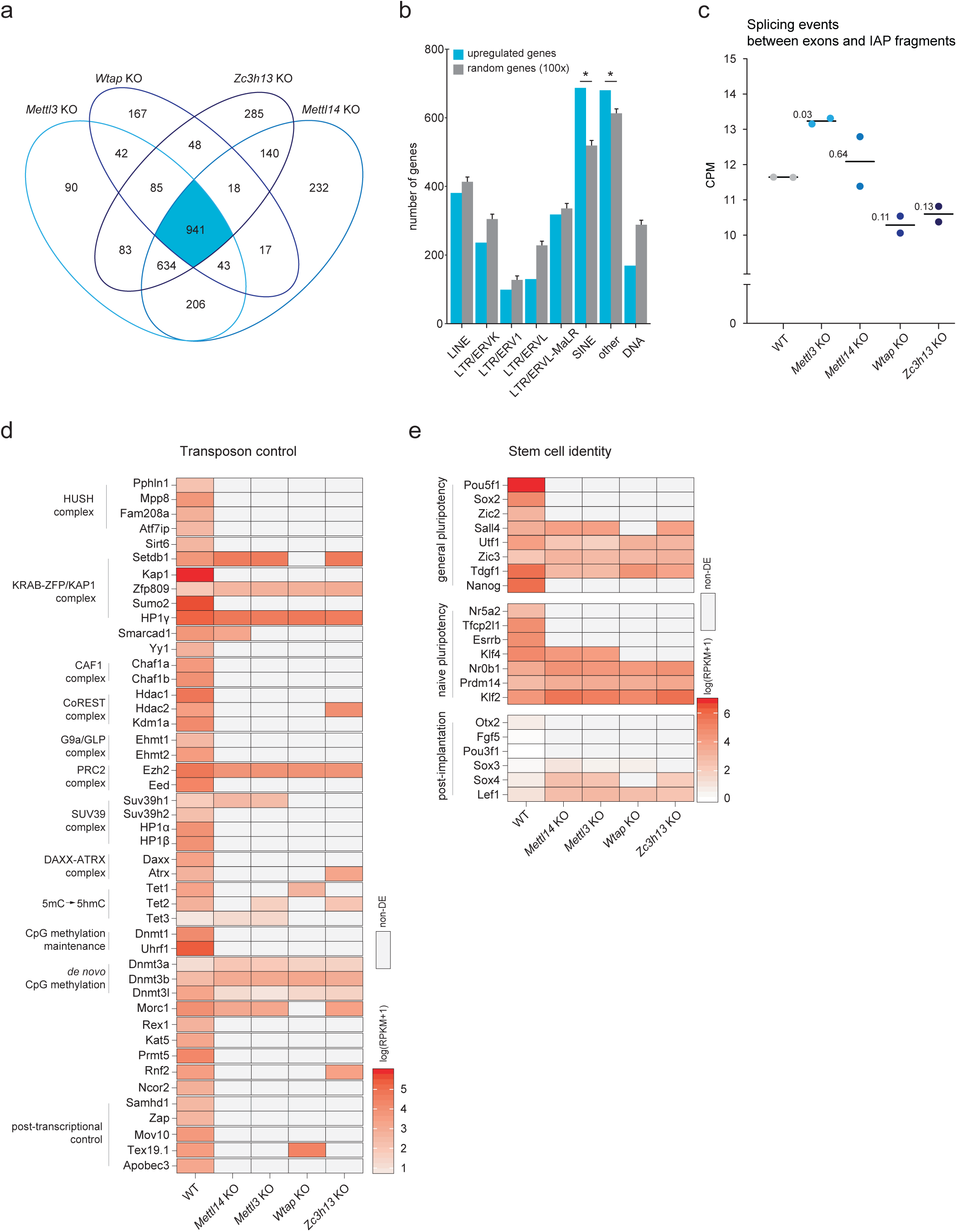
Analysis of gene expression in mutant ESCs of the m^6^A methyltransferase complex. **a**, Venn diagram for the upregulated genes (FDR<0.05 and log2FC>0.75) as assessed by RNA-seq in the indicated KO ESCs. **b**, Correlation between gene upregulation and proximity of retrotransposon annotations (−5kb to +1kb from the TSS). In blue are the 941 genes that are commonly upregulated in the four KO lines. Randomized control genes (100 randomizations) were annotated for comparison (grey). Permutation tests were performed. Asterisks indicate significant *P*-values (0.0099). **c**, Dot plot for normalized counts per million (CPM) of splicing events occurring between exons and RepeatMasker-annotated IAP elements in WT and indicated KOs. Numbers, which represent *P*-values for each condition, were assessed using Student’s t-test (KO *vs* WT); *n*=2 biological replicates). **d, e**, RNA-seq heatmaps show expression of selected retrotransposon regulators (**d**), and pluripotency and post-implantation markers (**e**) in WT and KO ESCs. Expression in KOs is only indicated when log2FC>0.75 and FDR<0.05; grey color (non-DE) reports no significant change in expression.

**Extended Data Fig. 6.**
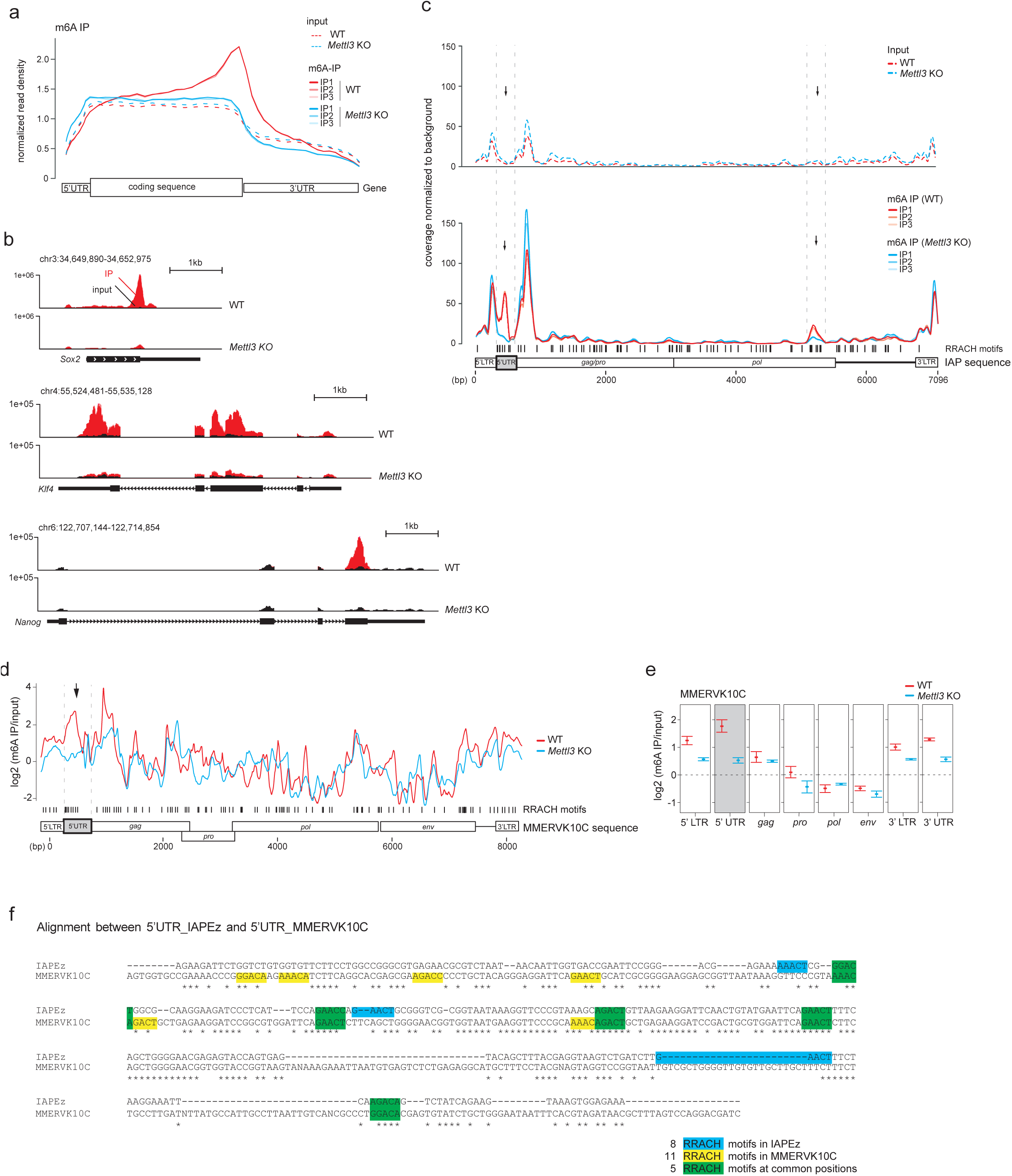
IAP mRNA undergoes METTL3-dependent m^6^A methylation. **a**, Normalized read density in WT (red) and *Mettl3-*KO (blue) across the 5’UTR, CDS, and 3’UTR of mRNA for the genes with at least one m^6^A peak in m^6^A RNA-IP (dashed lines represent respective inputs). **b**, m^6^A enrichment and distribution at representative genes known to undergo m^6^A methylation. Normalized read density (RPM) levels are shown as red shades: m^6^A-IP in WT and *Mettl3-*KO; black shades: m^6^A inputs. **c**, Background-normalized m^6^A signal coverage distribution across the IAPEz consensus sequence for three m^6^A-IP replicates in WT and *Mettl3*-KO (dashed lines represent respective inputs). Distribution of RRACH motif indicated as black lines. Black arrows indicate regions of m^6^A signal enrichments in WT that are lost upon *Mettl3* depletion. **d**, Average of input-normalized m^6^A signal intensities for three m^6^A-IP replicates along the MMERVK10C consensus sequence in WT (red) and *Mettl3-*KO (blue). Distribution of RRACH motifs is indicated as vertical black lines. The black arrow points to the 5’UTR region of METTL3-dependent m^6^A enrichment (present in WT and lost in *Mettl3-*KO). **e**, Average of m^6^A signal intensities for the indicated MMERVK10C sequence segments (m^6^A-IP replicates, *n*=3). **f**, Alignment between 5’UTRs of IAPEz and MMERVK10C consensus sequences. IAP-specific, MMERVK10C-specific and common RRACH motifs are indicated in blue, yellow and green, respectively.

**Extended Data Fig. 7.**
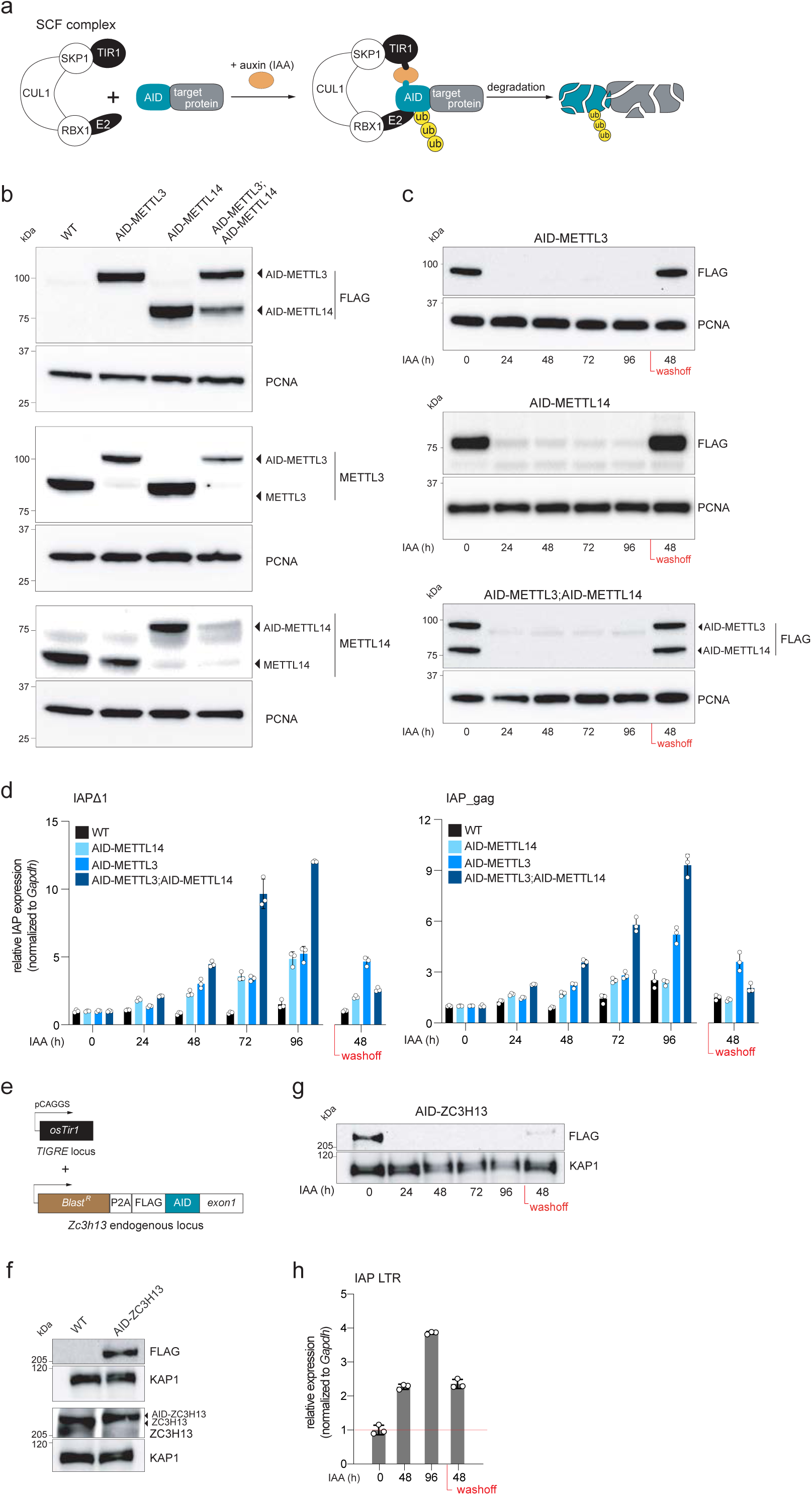
Auxin (IAA)-inducible degron of endogenous METTL3, METTL14 and ZC3H13. **a**, Schematic of TIR1 and SCF1 complex-dependent degradation of endogenously AID-tagged proteins in presence of IAA. **b**, Immunoblot demonstrating expression of endogenously 3xFLAG-AID-tagged METTL3 and METTL14 in single and double degron lines. TIR1 line was used as control for protein levels. PCNA serves as a loading control. **c**, Immunoblot demonstrating efficiency and reversibility of METTL3 and METTL14 depletion in 0-to-96h +IAA and 48h IAA washoff. PCNA serves as a loading control (related to Fig. 4d). **d**, RT-qPCR for IAP mRNA using Δ1- or *gag*-specific primers upon 0-96h of IAA treatment followed by 48h IAA removal in AID-METTL14 (light blue), AID-METTL3 (blue), AID-METTL3;AID-METTL14 dd (dark blue). TIR1 line was used as WT control (data shown as means ± s.d. from three technical replicates). IAP levels were normalized to *Gapdh* and expressed relative to levels at 0h (set to 1). **e**, Schematic of ZC3H13 degron ESC line. **f, g**, ZC3H13 auxin-dependent degron. Immunoblot demonstrating expression of endogenously 3xFLAG-AID-tagged ZC3H13 (**f**) and degron efficiency in presence of IAA (**g**). TIR1 line was used as control for protein levels. KAP1 serves as loading control. **h**, RT-qPCR analysis of IAP mRNA levels in ZC3H13 degron ESCs using LTR-specific primers in 0-48-96h of IAA treatment followed by 48h IAA washoff (data shown as means ± s.d. from three technical replicates). IAP levels were normalized to *Gapdh* and expressed relative to levels at 0h (set to 1).

**Extended Data Fig. 8.**
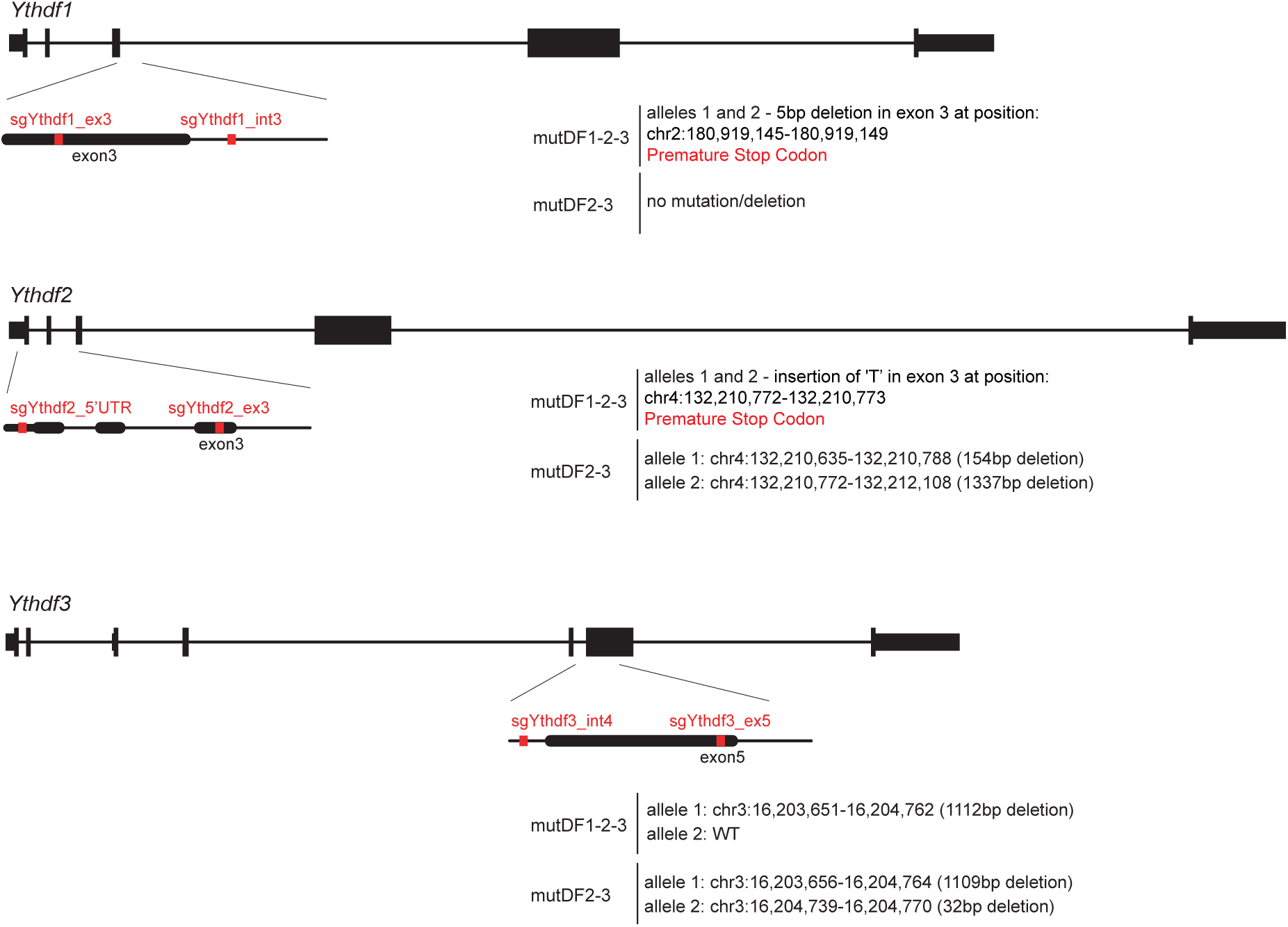
Generation of Y*thdf* mutant ESCs. Mutant **(mut)Ythdf2-3** and **mutYthdf1-2-3** cells were generated by CRISPR-Cas9 deletions. MutDF1-2-3 carried homozygous mutations in *Ythdf1* and *Ythdf2* and a heterozygous mutation in *Ythdf3*, and mutDF2-3 carried homozygous mutations at *Ythdf2* and *Ythdf3* genes. Schematic representation showing sgRNA sequences; mutation/deletion information based on Sanger sequencing is provided.

## Supplementary Information

### Supplementary Figures

Supplementary Figure 1 | Uncropped images of western blot gels

### Supplementary Tables

Supplementary Table 1: IAPEz nucleotide sequence used in IAPEz reporter cassette; primer sequences used for CRISPR-Cas9 library amplification and sequencing for screens I and II; primers used in RT-qPCR experiments.

Supplementary Table 2: List of sgRNA sequences used for generation of mutant and degron lines.

Supplementary Table 3: List of antibodies.

Supplementary Table 4: CRISPR-Cas9 knockout screen I and II results (MaGeCK)

Supplementary Table 5: Sequencing statistics for the CRISPR-Cas9 Screen I and II, RNA-seq and MeRIP-seq.

Supplementary Table 6: *Mettl3*-, *Mettl14*-, *Wtap*- and *Zc3h13*-KO RNA-seq results for differentially-expressed retrotransposon elements.

## Data availability

Data have been deposited in the Gene Expression Omnibus (GEO) under accession GSE145616.

## Acknowledgements

We would like to thank the members of the Bourc’his lab for their support. We are grateful to M. Greenberg and A. Shkumatava for critical reading of the manuscript, G. Cristofari for bioinformatic suggestions, I. Pinheiro for help with FACS analysis and E. Nora for vectors targeting *ROSA26* and *TIGRE* loci. Cell illustrations used in the Fig. 1 were obtained from https://smart.servier.com/ with slight changes. We acknowledge the ICGex NGS platform of the Institut Curie -supported by grants ANR-10-EQPX-03 (Equipex) and ANR-10-INBS-09-08 (France Génomique) from the Agence National de la Recherche- and the Cell and Tissue Imaging Platform (PICT-IBiSA) of Institut Curie -member of the French National Research Infrastructure France-BioImaging (ANR-10-INBS-04)”. The laboratory of D.B. is part of the Laboratory d’Excellence LABEX (LABEX) entitled DEEP (11-LBX0044). This work was supported by the Fondation Bettencourt Schueller, the Association Robert Debré pour la Recherche Médicale (ARDRM), the Fondation pour la Recherche Médicale (FRM) and the Association de Recherche contre le Cancer (ARC-PJA-20191209637). T.C. was a recipient of an EMBO postdoctoral fellowship and E.R. is supported by a PhD fellowship from la Ligue contre le Cancer.

## Author contribution

D.B. and T.C. conceived and designed the study. T.C. performed the genetic screen, genetic engineering of the different ESC lines (reporter, genetic knock-outs and auxin-degron), RNA-seq, MeRIP-seq, auxin-degron experiments, actinomycin D assays, m^6^A quantification by ELISA and immunoblots. E.R. contributed to generating and characterizing ESC knock-outs of *Mettl3, Mettl14, Wtap, Zc3h13, Ythfd1, Ythfd2* and *Ythfd3* and provided assistance with auxin-degron experiments. S.R. and C.F. provided assistance with the genetic screen, and S.L. and M.D. with MeRIP-seq. F.D. provided degron-targeting vectors and helped with degron design. A.T. performed the bioinformatic analyses. D.B. and T.C. interpreted the data and wrote the manuscript. All authors read and approved the final manuscript.

